# Genotyping of zebrafish embryos fixed by paraformaldehyde

**DOI:** 10.1101/118331

**Authors:** Xiaojie Yang, Qilin Gu

## Abstract

The rapid generation of various species and strains of zebrafish using CRISPR/Cas9 technology has dramatically accelerated the interrogation of gene function in *vivo*. So far, several approaches for genotyping of unfixed genome-modified animals have been successfully developed, such as Surveyor assays, T7 endonuclease 1 (T7E1) assays, High Resolution Melt Analysis (HRMA) and PAGE-based genotyping approach. However, there is few published genotyping protocol for the embryos of lethal zebrafish mutants fixed by paraformaldehyde. We have designed this genotyping protocol so that it can be performed in a single step with reliable results. The protocol covers all steps to obtain the genotypes.

## 1 Introduction

Since the 1960s, the zebrafish (Danio rerio) has become increasingly important to scientific research. It has many characteristics that make it a valuable model for studying human genetics and disease[1–4]. Zebrafish mutations provide an invaluable resource for dissecting genetic pathways that regulate vertebrate development, physiology, and behavior [5–12]. The optical clarity and accessibility of the zebrafish embryo allow exquisite analysis of mutant phenotype and a detailed understanding of gene function at the organismal and cellular level. Techniques have been developed for genotyping of unfixed indel mutants at the targeted site, such as Surveyor assays [13–15], T7 endonuclease 1 (T7E1) assays[16, 17], High Resolution Melt Analysis (HRMA) [18, 19]and PAGE-based genotyping approach [20].

Zebrafish embryos are nearly transparent which allows researchers to easily examine the development of internal structures, using many techniques to describe the expression pattern of developmentally regulated genes, such as whole-mount *in situ* hybridization (ISH)[3, 21], Whole-mount AP staining[22], whole-mount Immunofluorescence[4], whole-mount alcian blue staining[2]. All of these methods require fixing the embryos using paraformaldehyde. The only way to get the homozygotes of lethal mutants is incross the heterozygotes. However, the current publications show the phenotype of lethal mutants but not describing the genotyping methods. And also there is few publications describe how to extract DNA for paraformaldehyde-fixed zebrafish embryos. To our knowledge, this is the first report of an efficient and accurate method of genotyping zebrafish samples that have been fixed by paraformaldehyde. This method identifies homozygous mutants, heterozygotes, and homozygous wild type embryos.

## 2 Materials

### 2.1 Molecular Biology Reagents

1. Phosphate-Buffered Saline (PBS, pH 7.4)
2. Tween 20
3. PBST (1 × PBS, 0.1% (vol/vol) Tween 20)
4. KAPA Express Extract Kit (Kapa Biosystems, cat. no. KK7103)
5. Q5® High-Fidelity 2× Master Mix (NEB, cat. no. M0492L)
6. T7 Endonuclease I (NEB, cat. no. M0302L)
7. Gel Loading Dye
8. Agarose
9. GelRed Necleic Acid Stain (Phenix Research Products, cat. no. RGB-4103)
10. Tri-borate-EDTA (TBE Buffer)

### 2.2 Equipment

1. Low Speed Orbital Shaker
2. Heating block (75 °C, 95 °C)
3. 0.6-mL Microcentrifuge Tubes
4. 0.2-mL PCR tubes
5. PCR Thermocycler
6. DNA Electrophoresis Cells & Systems
7. Gel Imaging Systems

## 3 Methods

### 3.1 Preparation of embryos for DNA extraction

1. Collect the paraformaldehyde-fixed embryos in the 0.6-mL Microcentrifuge Tubes (one embryo per tube). Use one wild type embryo as negative control (NC).
2. Add 0.5 mL PBT to each tube, wash for three times, 5 min per wash on a low speed orbital shaker.
3. Add 0.5 mL 1× PBS to each tube, wash for three times, 5 min per wash on a low speed orbital shaker.
4. Wash the embryos in Mini-Q water for a final 5 min on a low speed orbital shaker.
5. Carefully aspirate the supernatant.
6. Spin at 10,000 g for 30 s, remove residual water with the 20 μL tips. The embryos are ready for lysis.

### 3.2 Extraction of DNA for PCR

1. After thawing reagents, vorter to mix and spin briefly, keep all reagents on ice until use.
2. Prepare the KAPA lysis buffer tabulated below on ice.

3. Add 50 μL KAPA lysis buffer to each 0.6-mL tube containing zebrafish embryo.
4. Perform lysis in a heating block using the conditions tabulated below (*see* **Note 1**).

| Component | Amount per reaction | Final |
| --- | --- | --- |
| 10× KAPA Express Extract Buffer | 5 μL | 1× |
| 1 U μL<sup>-1</sup> KAPA Express Extract Enzyme | 1 μL | 1 U |
| PCR-grade water | 44 μL | - |

5. After lysis, spin at 10,000 g for 30 s. For alternatives *see* **Note 2.**

3. Add 50 μL KAPA lysis buffer to each 0.6-mL tube containing zebrafish embryo. 4. Perform lysis in a heating block using the conditions tabulated below (*see Note 1*).
| Step | Temperature | Duration |
| --- | --- | --- |
| Lysis | 75 $^{\circ}$C | 15 min |
| Enzyme inactivation | 95 $^{\circ}$C | 5 min |

### 3.3 PCR amplification

1. After thawing reagents, vorter to mix and spin briefly, keep all reagents on ice until use.
2. Set up a separate 20 μL PCR in a 0.2-mL sterile tube, as tabulated below (see **Notes 3-5**).

3. Gently flick the Master Mix tube to mix and spin briefly.
4. Dispense 19 μL of Master Mix into 0.2-mL PCR tubes for each sample to be amplified.
5. Add 1 μL of DNA samples to each corresponding PCR tube.
6. Gently flick PCR tubes to mix the DNA and PCR Master Mix.
7. Run the PCR using the conditions tabulated below.

| Component | Amount per reaction | Final |
| --- | --- | --- |
| Forward primer (10 μM) | 0.4 μL | 0.2 μM |
| Reverse primer (10 μM) | 0.4 μL | 0.2 μM |
| Q5<sup>®</sup> High-Fidelity Master Mix (2×) | 10 μL | 1× |
| Template DNA | 1 μL | - |
| Sterile water | 8.2 μL | - |

8. Cool the amplified PCR product to 4 °C and store at −20 °C for weeks, or continue to next step.

| Cycle number | Denature | Anneal and Extend |
| --- | --- | --- |
| 1 | 95 °C, 3 min | - |
| 2-36 | 98 °C, 10 s | 68 °C, 30 s |
| 37 | - | 72 °C, 5 min |

### 3.4 SURVERYOR and T7 endonuclease I cleavage

1. To perform SURVERYOR assays, aliquot 5 μL PCR products in a 0.2-mL sterile tube, place PCR tubes in thermal cycler and run the re-annealing program to enable heteroduplex formation using the conditions tabulated below.

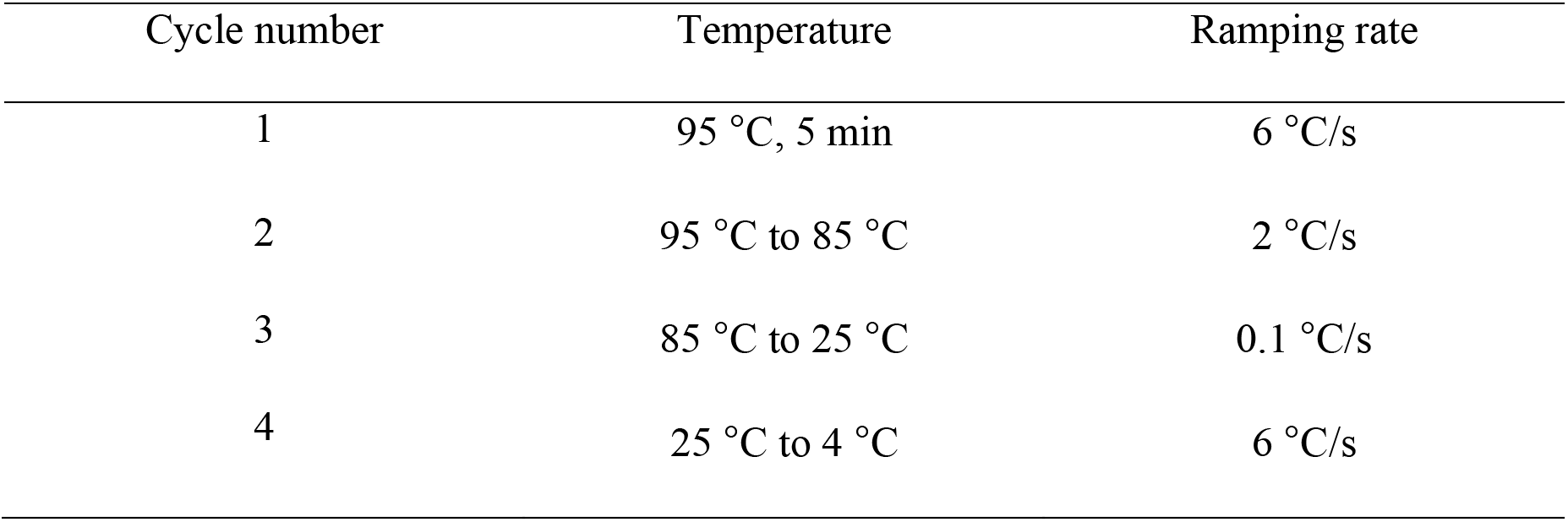
2. Cool the amplified PCR product to 4 μC and store at −20 μC for weeks, or continue to next step.
3. Aliquot 2 μL re-annealing PCR product in a 0.2-mL sterile tube. Set up a separate 20 μL T7 Endonuclease I cleaving reactions as tabulated below.

| Cycle number | Temperature | Ramping rate |
| --- | --- | --- |
| 1 | 95 °C, 5 min | 6 °C/s |
| 2 | 95 °C to 85 °C | 2 °C/s |
| 3 | 85 °C to 25 °C | 0.1 °C/s |
| 4 | 25 °C to 4 °C | 6 °C/s |

4. Gently flick the tube to mix the T7 Endonuclease I cleaving buffer and spin briefly.
5. Dispense 18 μL of cleaving mix into 0.2-mL PCR tubes for each sample.
6. Add 2 μL of re-annealing product to each corresponding PCR tube.
7. Gently flick PCR tubes to mix the DNA and cleaving mix.
8. Place at 37 °C for 2 h. The cleaving product can be stored at −20 °C for weeks.

| Component | Amount per reaction | Final |
| --- | --- | --- |
| Re-annealing PCR product | 2 µL | - |
| NEBuffer 2 (10×) | 2 µL | 1× |
| T7 Endonuclease I (10 U µL<sup>-1</sup>) | 0.1 µL | 1 U |
| Water | 15.9 µL | - |

### 3.5 Agarose Gel Electrophoresis and Imaging

1. Prepare a 1.5% (g/ml) agarose gel using TBE buffer (*see* **Note 6**).
2. Add 3 μL loading dye to each reaction, mix and spin briefly.
3. Electrophorese the total product.
4. Run the gel, 120 volts for 20 min.
5. Document the gel image to interpret genotype.
6. The heterozygotes can be distinguished from the electrophoretic results that show three bands. The wild type and homozygotes will only show one band and need be genotyping again.

### 3.6 Genotyping the homozygotes

1. Separate the PCR product from **step 8** (subheading 3.3) that show one band in **step 6** (subheading 3.5).
2. Aliquot 2.5 μL PCR products in a 0.2-mL sterile tube, mix with 2.5 μL NC PCR products.
3. Repeat the **steps** from subheading 3.4 to 3.5.
4. The electrophoretic results that show one or three bands are wild type or homozygotes, respectively.

### Anticipated Results

If the target site contains a restriction endonuclease site near the PAM region, it’s optional to use the restriction endonuclease to cleave the PCR products. The WT samples will be completely cut into two bands; the heterozygotes will partly be cleaved into two bands and partly be uncut; and the homozygotes will show one uncut band (Fig. 1A and 1B; 2A and 2B). If the target site contains no restriction endonuclease site, the PCR products will subject to SURVERYOR assays. The re-annealing products of the heterozygotes will partly form heteroduplex and be cleaved by T7 Endonuclease I into two bands. At this point, the WT and homozygotes will show one band; the heterozygotes will show three bands (Fig. 1C and 2C). We need separate the one-band products and repeat the SURVERYOR assays. By mixing the one-band products with the negative control (Wild type sample), the homozygous samples will partly form heteroduplex be cleaved into two bands. While the WT samples will only show one band (Fig. 1D and 2D).

**Figure 1.**
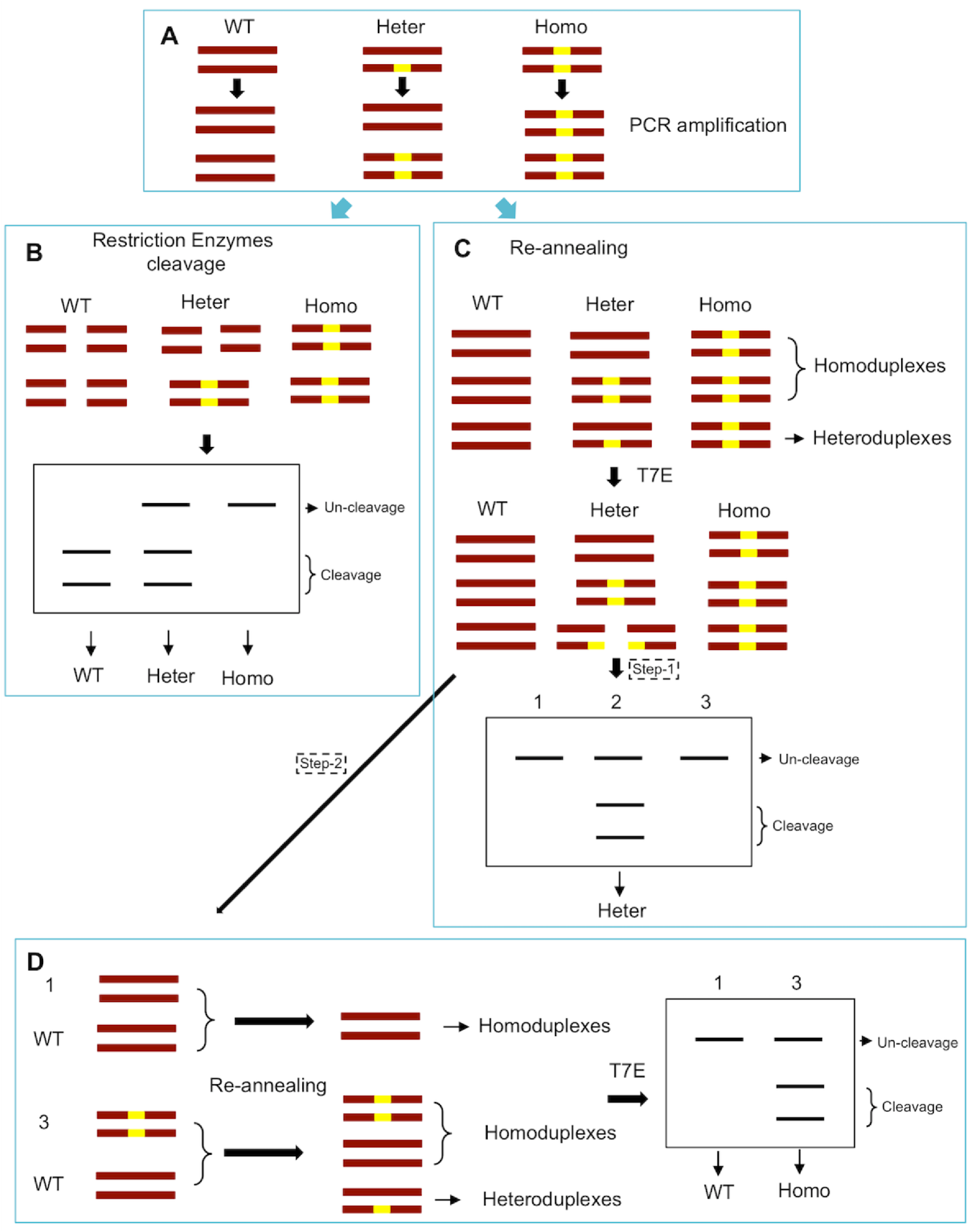
Schematic overview of T7E I-based genotyping protocol for identification of CRISPR/Cas9-mediated mutations. (**A**) Illustration of DNA formation in different genotype samples during PCR amplification. Dark red bars represent DNA strands in cells harboring monoallelic mutations (yellow box). WT, wild type; Heter, heterozygous; Homo, homozygous. (**B**) The target site contains a restriction endonuclease site, then use the restriction endonuclease to cleave the PCR products. The WT samples will be completely cut into two bands; the heterozygotes will partly be cleaved into two bands and partly be uncut; and the homozygotes will show one uncut band. (**C**) The target site contains no restriction endonuclease site, the PCR products will subject to SURVERYOR assays. The re-annealing products of the heterozygotes will partly form heteroduplex and be cleaved by T7 Endonuclease I (T7E) into two bands. At this point, the WT and homozygotes will show one band; the heterozygotes will show three bands. (**D**) The one-band products in C were separated and repeated the SURVERYOR assays. By mixing the one-band products with the negative control (Wild type sample), the homozygous samples will partly form heteroduplex be cleaved into two bands. While the WT samples will only show one band.

**Figure 2.**
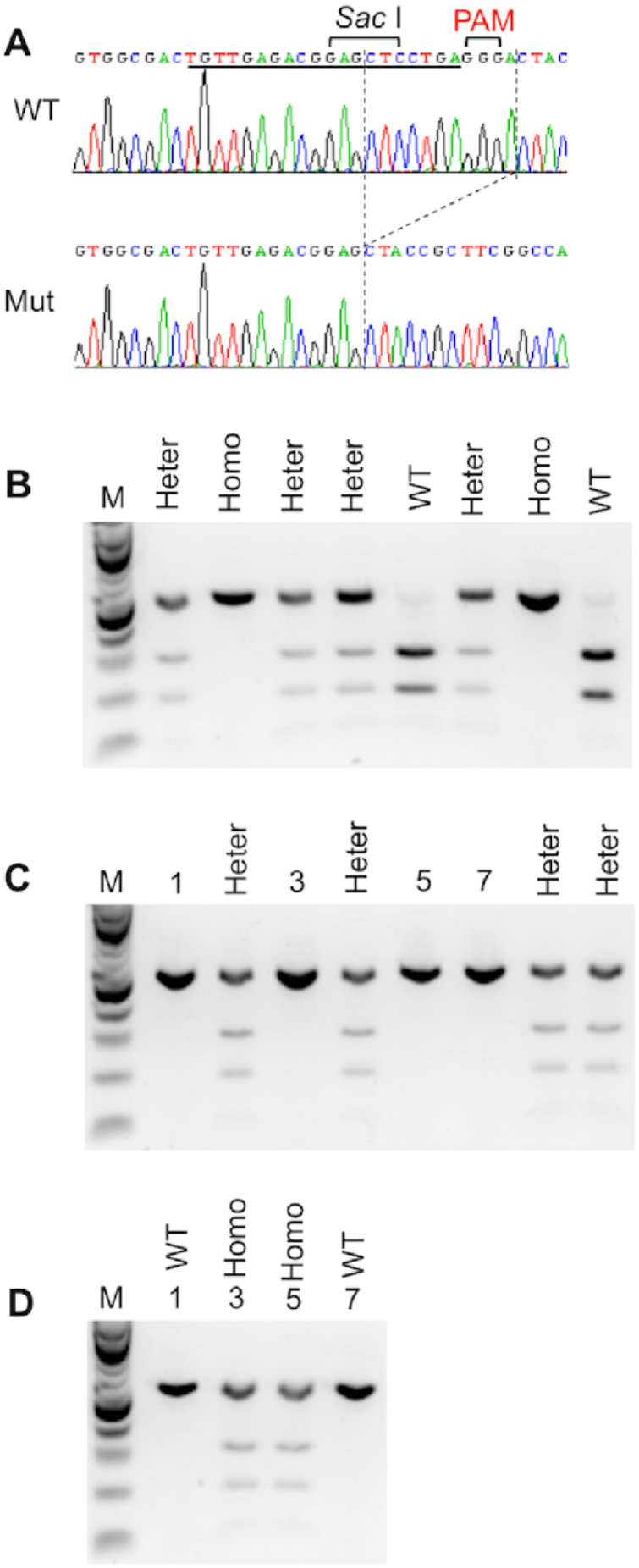
Detection of CRISPR/Cas9-mediated genome-modified zebrafish mutant by T7E I-based genotyping assay. (**A**) The target site contains a restriction endonuclease site (*Sac* I) in the wild type genome. WT, wild type; Mut, mutant. (**B**) The WT samples are completely cut into two bands by *Sac* I digestion; the heterozygotes are partly cleaved into two bands and partly be uncut; and the homozygotes show one uncut band. WT, wild type; Heter, heterozygous; Homo, homozygous. (**C**) The PCR products will subject to SURVERYOR assays. The WT and homozygotes will show one band (line 1, 3, 5 and 7); the heterozygotes will show three bands. (**D**) The one-band products (line 1, 3, 5 and 7) were separated and repeated the SURVERYOR assays. By mixing the one-band products with the negative control (Wild type sample), the homozygous samples are cleaved into two bands (line 3 and 5). While the WT samples only show one band (line 1 and 7).

## 4 Notes

1. Mix occasionally until no intact embryos visible in the tube. Unlike digestion with proteinase K, KAPA Express Extract does not completely degrade the tissue. This does not have a negative impact on the downstream PCR.
2. Extract solutions are stable at 4 °C for weeks or store at −20 °C for at least 6 months.
3. Reaction volumes >20 μL are not recommended, as reaction efficiency may be compromised.
4. Other PCR Mix contains loading dye is not recommended, because the PCR product might not be directly used as template for the following T7 Endonuclease I cleaving reactions.
5. Be careful not to contaminate the PCRs. Use sterile tubes and filter tips and wear gloves.
6. We recommend to using 1× GelRed to replace the highly toxic ethidium bromide.

## Acknowledgements

X.Y. is supported by the postdoctoral fellowship from the American Heart Association.

## Author Contributions

X.Y. and Q.G. conceived, designed and performed the experiments, analyzed the data, wrote the paper, prepared figures, reviewed drafts of the paper.

